# Structural Variation and Polysaccharide Profiling of Intervessel Pit Membranes

**DOI:** 10.1101/2021.06.07.447397

**Authors:** Qiang Sun

## Abstract

Functional roles of intervessel pit membrane (PM) depend on its structure and polysaccharide composition, which are mostly unknown or lack of accurate information. This study uses grapevine as a model plant and an immunogold-scanning electron microscopy technique to simultaneously analyze structures and polysaccharide compositions of intervessel PMs in relation to their functions. Intervessel PMs with different structural integrity were found in functional xylem with about 90 % of them being intact with a smooth or relatively smooth surface and the rest 10 % with progressively degraded structures. The results also elucidated details of the removal process of wall materials from surface toward its depth during the natural intervessel PM degradation. Four groups of pectic and hemicellulosic polysaccharides were present in intervessel PMs but displayed differential spatial distributions and quantities: weakly methyl-esterified homogalacturonans abundant in the surficial layers, heavily methyl-esterified homogalacturonans and xylans mostly in deep layers, and fucosylated xyloglucans relatively uniform in presence at different depths of an intervessel PM. This information is crucial to reveal the polysaccharide profiling of primary cell wall and to understand intervessel PM’s roles in the safety and regulation of water transport as well as the plant susceptibility to vascular diseases.

## Introduction

As a major type of water conducting tissue, vessel is a hollow, both-end sealed tube which is itself composed of many axially end-to-end connected vessel elements. Individual vessels are limited in length and overlap in an organ’s axial direction, forming a network in the xylem throughout a plant body (Mauseth, 1988; Evert, 2006). At each overlapping region of the network, contiguous vessels have a direct lateral wall contact where pit pairs (PPs) are present. Since secondary wall deposition does not occur at PPs, two contiguous vessels at each PP are separated only by the two vessel elements’ primary cell walls and one middle lamella, which are collectively called intervessel pit membrane (PM, Esau, 1977).

Intervessel PM is an important structure that affects water transport through the vessel system in a plant and has therefore attracted a lot of research attention (e.g., Tyree and Sperry, 1989; Tyree and Zimmermann, 2002; Stroock et al., 2014). Early studies focused on the structure of intervessel PMs. Since intervessel PMs are not covered by secondary cell walls, they are subject to the attack of hydrolytic enzymes from the tonoplast breakdown during the vessel element differentiation. The intervessel PMs’ non-cellulosic components are presumably removed via the hydrolysis to increase their porosities (Schmid and Machado, 1968; O’brien, 1970, 1974; Butterfield and Meylan, 1982). This modification is thought to facilitate the across-vessel movement of water and dissolved minerals, consequently contributing to the effective axial water transport through the vessel network (Esau, 1977; Tyree and Zimmermann, 2002). However, information is still lacking in terms of details and/or direct evidence of how intervessel PMs are structurally modified, what kinds of polysaccharides exist and how they are removed from the intervessel PMs during this modification process.

Another research focus of intervessel PM has been on its impacts on the safety of water transport in a vessel network. Although intervessel PMs are adapted to transport water in the xylem due to lack of secondary wall deposition, they were found to still account for 50-90 % of the hydraulic resistance in a plant, depending on species (Wheeler et al., 2005; Choat et al., 2006; Hacke et al., 2006). It has been suggested that this hydraulic resistance should be due to extremely small pore sizes (5-20 nm) of intervessel PMs (Choat et al., 2003; Pérez-Donoso et al., 2010). On the other hand, the small porosity of intervessel PMs is believed to help prevent air bubbles in some embolized vessels (dysfunctional vessels) from spreading to adjacent vessels in the vessel network. This consequently helps maintain continuous water columns in the vessel system, contributing to the safety of water conduction in the xylem (Sperry and Tyree, 1988; Choat et al., 2005; Rockwell et al., 2014). Therefore, accurate information about the structures, especially porosity, of intervessel PMs is crucial to better understand how xylem tissue/its vessel system may maintain the safety for water transport. However, data on this aspect are not yet enough for accurate analyses.

Intervessel PM itself is believed to potentially interact with solutes transported in the vessel system, directly regulating the water flow rate in the xylem (Zwieniecki et al., 2001; van Doorn et al., 2011). It has been found that ion concentration in xylem sap has impacts on the hydraulic conductivity of the xylem. Increased hydraulic conductivities were demonstrated in many species when the ion concentrations were above certain levels (Zimmermann, 1978; van Ieperen et al., 2000; López-Portillo et al., 2005; Domec et al., 2007; Gascó et al., 2006; Nardini et al., 2007; Cochard et al., 2010; Gortan et al., 2011). Currently, two major hypotheses are available to explain this phenomenon (Zwieniecki et al., 2001; van Doorn et al., 2011). Although both attribute the ion-increased hydraulic conductivity to ion interactions with polysaccharides of intervessel PMs, the polysaccharides involved belong to different groups, i.e., pectins (Zwieniecki et al., 2001) or any polyelectrolyte polymers but not pectins (van Doorn et al., 2011). There are still a lot of controversies about the absence/presence of pectic polysaccharides in mature intervessel PMs, which underlines the two hypotheses. Therefore, accurate information about the polysaccharide composition of intervessel PM will be a key to understand any direct intervessel PM’s role in regulating the water flow rate in xylem.

Intervessel PM is also a target in the study of plant vascular diseases. Many vascular diseases are caused by xylem-limited pathogens, and their symptom development is largely dependent upon the systemic spread of pathogen cells in a host plant through its vessel system (Purcell and Hopkins, 1996; Bove and Garnier, 2002; Sicard et al., 2018). Intervessel PMs are believed to be barriers for this spread due to their small porosity; that is, pores in intervessel PMs are much smaller than bacterial pathogens (Pérez-Donoso et al., 2010; Sun et al., 2011). Some analyses have strongly suggested that pathogens secrete cell wall-degrading enzymes (CWDEs) to remove certain polysaccharides from intervessel PMs, enlarging the PMs’ porosity for the pathogen spread in the vessel system (Simpson et al., 2000; Roper et al., 2007; Sun et al., 2011; Ingel et al., 2019). However, little is known in terms of detailed interactions between the CWDEs and intervessel PMs. Therefore, understanding the polysaccharide composition and spatial distribution in intervessel PMs is essential to analyze the pathogen-host plant interactions, consequently elucidating disease susceptibility mechanisms of host plants.

The *in-situ* detection of pectic and hemicellulosic polysaccharides in intervessel PMs is also a research focus and has been based mostly on light microscopy (LM) with dye staining methods (Catesson, 1983; Jansen et al., 2004; Arend et al., 2008) and, in few cases, on transmission electron microscopy (TEM) with immunocytochemical technique (Plavcová and Hacke, 2011; Kim and Daniel, 2013). These techniques have revealed important information about cell wall chemical composition but showed major limitations when comprehensive analyses on the structures and polysaccharide compositions of intervessel PMs are needed (to be described in detail in the Discussion). In this study, we used an immunogold-scanning electron microscopy (SEM) protocol (Sun et al., 2017), which combines an immunocytochemical technique with SEM. The SEM can reveal, at different degrees of resolution, surface structures (intact surface and exposed underlayers) of intervessel PMs as well as those on sections. This immunogold-SEM protocol has been proved powerful in exploring the structures, polysaccharide compositions and functions of cell walls all together (Sun et al., 2017). With an expanding list of monoclonal antibodies (mAbs) and carbohydrate-binding modules (CBMs) for cell wall polysaccharide detection, this technique offers an excellent approach to visualize a large variety of polysaccharides (Willats et al., 2001; McCartney et al., 2004; Pattathil et al., 2010). The current study used this technique with grapevine as a model plant to simultaneously elucidate the structures and polysaccharide compositions of intervessel PMs.

## Results

### Structural features of intervessel PPs and PMs

Secondary xylem of grapevine stems contained four major types of cells: vessel element, fiber, axial parenchyma cell and ray parenchyma cell (Figure 1). A vessel made up of axially end-on-end connected vessel elements may have a direct cell wall contact with the four types of cells and formed PPs in the contact wall with all the cell types except fiber (Figure 1, A). PPs between two contiguous vessels (a.k.a intervessel PPs) occurred in a tight scalariform (ladder-like) pattern axially along the vessels’ contact wall and each PP is stretched horizontally throughout the width of the contact wall (Figure 1, A and B). Intervessel PPs were bordered PPs (Figure 1, A and C). When secondary wall borders of an intervessel pit are in place, only a small portion of its intervessel PM is visible from its surface view (Figure 1, B and D). The whole intervessel PM can be revealed only after the secondary wall borders of a pit (Figure 1, C and D) are removed (Figure 2).

**Figure 1.**
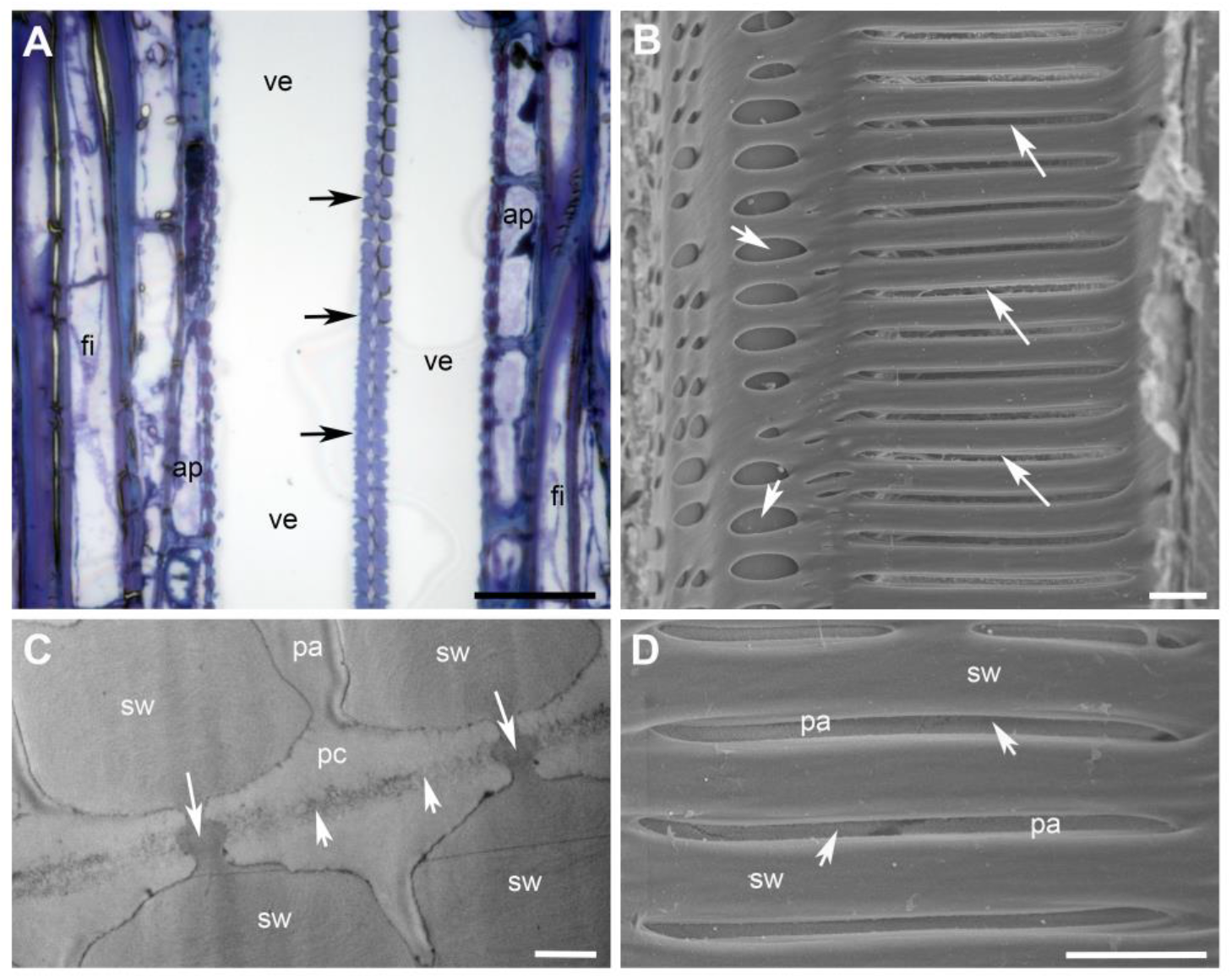
Morphology of intervessel pit pairs (PPs) and pit membranes (PMs) in secondary xylem of grapevine stems. A, Light micrograph of secondary xylem in a tangential longitudinal section, showing an array of densely arranged intervessel PPs (arrows) at the sectional view of the contact wall of two contiguous vessels (ve). Also shown in the image are axial parenchyma cells (ap) and xylem fibers (fi). B and D, SEM images of intervessel PPs at the surface view of the lateral wall of a cut-open vessel. B, Intervessel pits (long arrows) are scalariformly arranged in the axial direction of a vessel element and vessel-parenchyma pits (short arrows) are oval or transversely elongated. C, TEM image of an intervessel PP at a sectional view, showing that the PP contains two opposing bordered pits separated by a thin PM (short arrows). Secondary wall borders (sw) of each pit cover narrow portions (long arrows) of the vessel’s primary cell wall, enclose a large pit chamber (pc), and leaves a narrow pit aperture (pa). D, An enlarged image of some intervessel pits, showing that each pit has a narrow pit aperture slit (pa) flanked by secondary wall borders (sw) and that a small portion of intervessel PM (arrows) is visible through each pit aperture slit. Scale bar equals 40 μm in A, 10 μm in B and D, and 1 μm in C.

**Figure 2.**
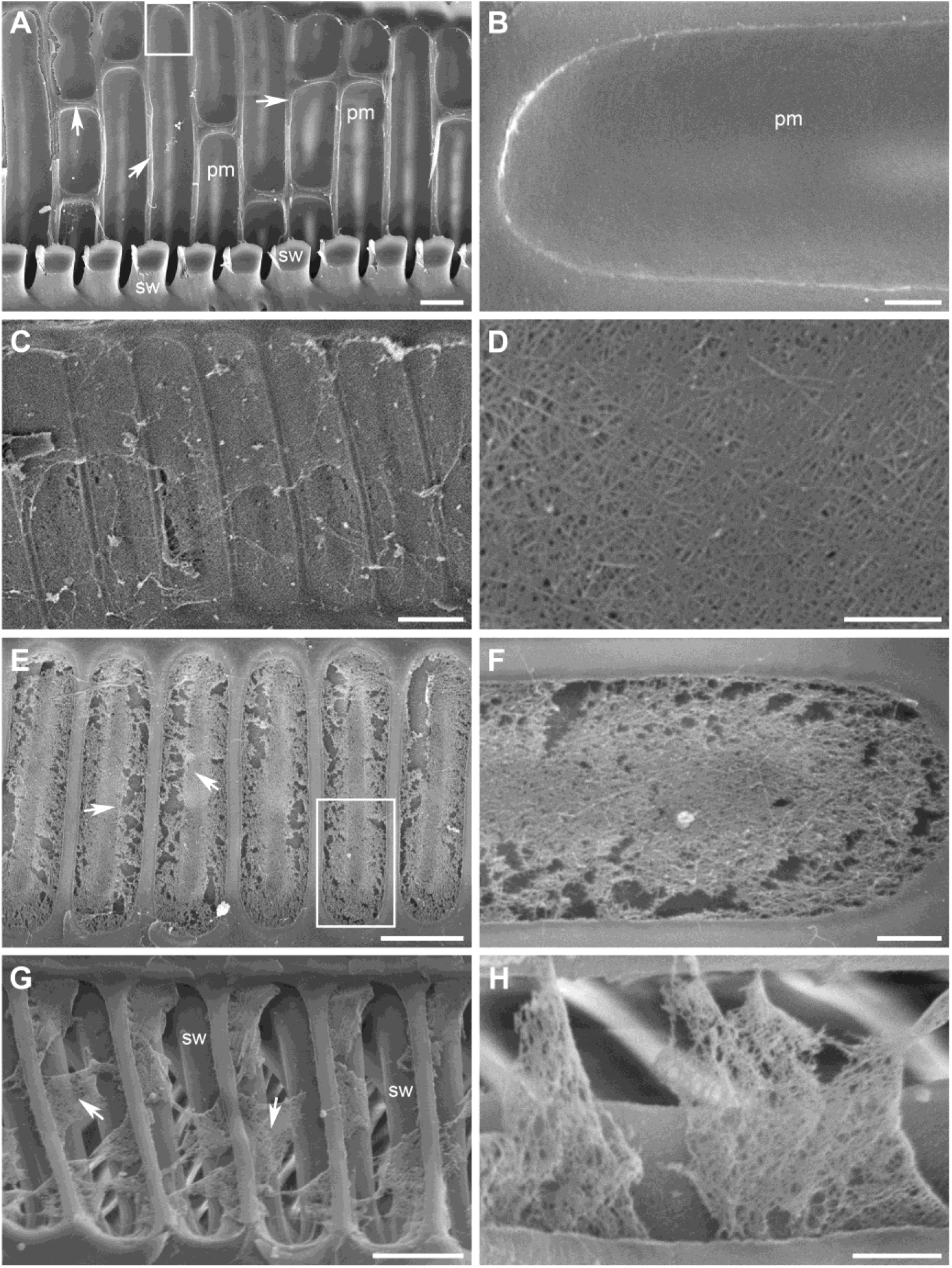
Representative SEM images demonstrating surface structural variations of intervessel PMs in secondary xylem of grapevine stems. A and B, Type I Intervessel PMs. A, Intervessel PMs (pm) are exposed, after removal of most of the secondary wall borders (sw), on the lateral wall of a longitudinally cut-open vessel element. Also exposed are strip primary wall regions (arrows) that were originally laid underneath the secondary wall and separate adjacent intervessel PMs. B, An enlarged image of the framed region in (A) with a 90° counterclockwise rotation, showing that the intervessel PM (pm) is intact and has a very smooth surface without a visible microfibril network. C and D, Type II intervessel PMs. C, Intervessel PMs are intact mostly with a smooth surface but have portions with a less smooth surface. D, Enlargement of a Type II intervessel PM, showing that randomly oriented microfibrils are exposed and small pores or indents are visible in some regions of the PM. E and F, Type III intervessel PMs. E, Increased porosity in intervessel PMs after a further loss of the cell wall material from the PMs. More wall material loss in the peripheral regions than in the central strip region (arrows) of each PM often resulted in large pores in the peripheral regions. F, An enlarged image of the framed region in (E) with a 90° counterclockwise rotation. The peripheral region has an exposed looser microfibril network with large irregular pores of different sizes while the central strip region has a relatively dense microfibril network with some filling wall material and smaller pores. G and H, Type IV intervessel PMs. G, Some portions of each intervessel PM have disappeared from their original places, forming large pores and the PM patches remaining in place (arrows) are very porous. The significant loss of intervessel PM integrity has made it visible the inner surface of the secondary wall borders (sw) of opposing pits on the vessel element underneath. H, An enlarged image of the intervessel PM regions remaining in place, showing the very porous microfibril networks with little amorphous material filled in. Scale bar equals 5 μm in A, C, E and G, and 1 μm in B, D, F and H.

Dimension of intervessel PM varied among intervessel PPs with a greater variation in width (21.7 ± 9.1 μm [mean ± sd, the same below], 5.6 – 45.8 μm in range, n = 246) than in height (5.1 ± 0.6 μm, 2.8 – 6.4 μm in range, n = 246). The width of an intervessel PM depended on the width of the contact cell wall where the PM was present at its two contiguous vessels. A same intervessel PM may have a slight difference in thickness in different regions when viewed under TEM (Figure 1, C). The thickest region of an intervessel PM was measured to represent its thickness. Intervessel PMs had the thickness of 636.6 ± 76.9 nm (509.1 – 781.8 nm in range, n = 20). Vessel-parenchyma PPs (those occurring between a vessel and a parenchyma cell) were oval shaped or transversely elongated and differed from intervessel PPs in size and arrangement pattern (Figure 1, B).

### Variations in the surface structural features of intervessel PMs

Surface features of intervessel PMs are one focus of this study due to their impacts on many functions of intervessel PMs. Intervessel PMs in the secondary xylem of grapevine stems showed diverse surface structural variations that were used to classify intervessel PMs to four distinguishable types (Figure 2). The four types corresponded to progressive stages in the integrity loss of an intervessel PM derived from the removal of its cell wall material.

Type I intervessel PMs were the predominant type out of the four. They had a homogeneously smooth surface over the whole PM region. Microfibrils and pores were not distinguishable from the PMs even at a magnification of up to 30,000 (Figure 2, A and B). These surface features have indicated presence of amorphous wall material covering microfibrils on the surface of intervessel PMs. This intervessel PM type represents no or the least damage on the structural integrity of intervessel PMs.

Another common type of intervessel PM (Type II) had a large, relatively smooth surface region and some small, less smooth regions (Figure 2, C and D). The smooth region was present in a PM’s wide central stripe region, while the less smooth regions occurred in its very narrow peripheral region but were sometimes seen towards one side of its width (Figure 2, C). Different from that of Type I intervessel PMs, the large smooth region of a Type II intervessel PM usually revealed a microfibril network, in which microfibrils became largely distinguishable as dense randomly oriented, more-or-less straight threads and were filled with amorphous material in between. The network also exhibited sparsely distributed pores or dents of less than 50 nm in diameter (Figure 2, D). These surface features resulted from a removal of some superficial material from the PM and the pores/dents indicated where a more localized loss of the amorphous substance occurred. The regions with a less smooth surface also had pores, which were mostly circular or oval shaped with a diameter of up to 80 nm (few could be over 110 nm) and were dispersed in the network. These pores occurred at a higher density than those in the smooth central PM region. The presence of the relatively smooth and less smooth regions in a Type II intervessel PM indicated that the removal of a superficial layer from an intervessel PM did not occur in a uniform manner over the whole PM region.

In Type III intervessel PMs (Figure 2, E and F), the non-uniformity in the PM surface features increased significantly. Most commonly, the PM’s central strip region with a relatively dense microfibril network structure reduced in area and became rough in its surface, while the PM’s peripheral region appeared as a more porous structure. In the reduced central region, microfibrils were still distinguishable in a dense network with some associated amorphous material. But in the peripheral region, amorphous material associated with the microfibril network was further reduced in quantity. Pores were present in both regions. They were smaller (usually less than 150 nm) and mostly in oval or circular shapes in the central region, but much larger (usually up to 1.2 μm and some over 2.5 μm) and mostly irregular in shape in the peripheral region. The larger pores should have resulted from a localized loss of more amorphous material and some microfibrils. These surface features revealed that a progressive disintegration has occurred to intervessel PMs and that the removal of wall material resulted in a more loss in the structural integrity in the peripheral region than in the central region of an intervessel PM.

Type IV intervessel PMs showed significant structural differences from Type III ones. Type IV intervessel PMs have lost their integrity with some regions losing PM structure and revealing the secondary wall borders of the opposing pits of the vessel that was originally concealed underneath the PM (Figure 2, G and H). Few PM patches still in place in each PP were different in size and shape and appeared to be much looser networks after a loss of more amorphous material and microfibrils. Microfibrils remaining in each network were distinguishable with little amorphous structures associated and pores in the network were irregular in shape and usually over 100 nm in size. There were intervessel PPs where intervessel PMs were completely absent. The dissociation and highly increased porosity of Type IV intervessel PMs resulted from the further removal of amorphous material and microfibrils from a Type III intervessel PM and a subsequent partial or complete removal of the PM due to its much-weakened structures.

Although the four types of intervessel PMs were observed in a same secondary xylem tissue, a single vessel usually exhibited one type of intervessel PMs along its axial direction. However, a single vessel sometimes had two intervessel PM types that are at the consecutive PM degradation stages. The two types might be in a same vessel element or different vessel elements. In rare cases, structural features of two intervessel PM types at the consecutive stages may occur in different regions of a same intervessel PM (Figure 2, C).

Frequencies of the four types of intervessel PMs in secondary xylem were different (Figure 3), with Types I and II having a majority of presence (53.9 % and 36.8 %, respectively) and Type III an occurrence of 5.8 %. Type IV intervessel PMs were present at a very low frequency of 3.6 %.

**Figure 3.**
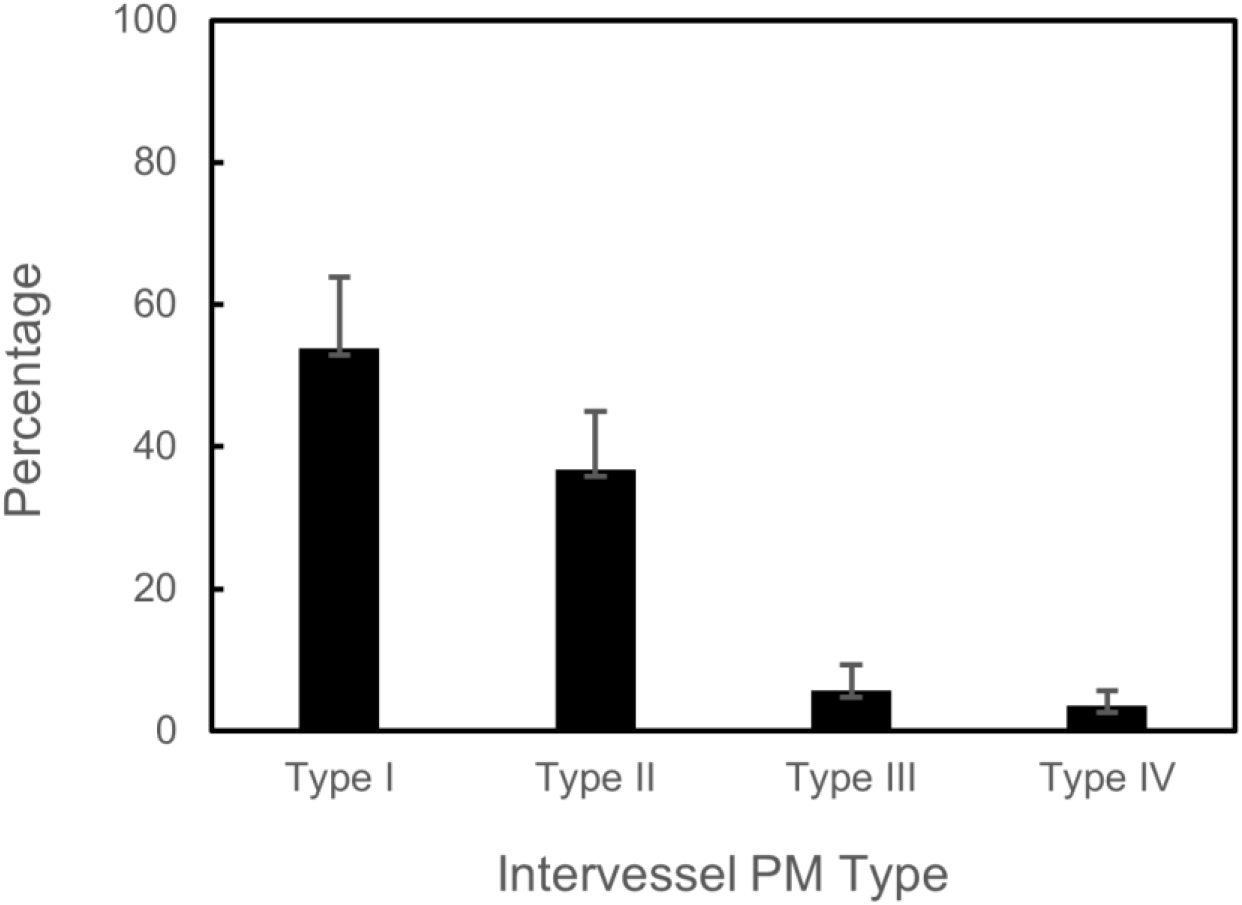
Frequencies of the four types of intervessel PMs in secondary xylem of grapevine stems. Each frequency is presented as mean with standard deviation and was based on 54 xylem specimens from 5 vines (8-13 specimens per vine).

### Spatial distributions and quantities of hemicellulosic and pectic polysaccharides in intervessel PMs

Intervessel PMs from Type I to Type IV represent different stages of the structural degradation of an intervessel PM and expose cell wall layers/materials at a PM’s different depths. When combined with the immunogold-SEM technique, they are ideal for exploring the spatial distribution and relative quantities of cell wall polysaccharides in an intervessel PM.

Immunolabelling the four types of intervessel PMs with four kinds of cell wall mAbs was used to detect two groups of hemicellulosic polysaccharides (xylans and fucosylated xyloglucans [F-XyGs]) and two groups of pectic polysaccharides (weakly methyl-esterified homogalacturonans [WMe-HGs] and heavily methyl-esterified homogalacturonans [HMe-HGs]), respectively in intervessel PMs. The information attained was then used to reveal the spatial organization of these major polysaccharides in an intervessel PM.

In all the three types of negative controls for the immunogold labeling experiments with each mAb (Figures 4, A and 6, A), no or very few randomly distributed bright particles were observed, indicating that the immunolabeling protocol in this study was adequate in suppressing any background noise derived from unspecific binding of mAb and/or secondary Ab. Therefore, the presence/absence of bright particles observed in each immunolabeling experiment with a particular mAb and its corresponding secondary Ab truly revealed the presence/absence of its particular polysaccharides. Accordingly, densities of the bright particles among different PM types for each mAb treatment reveal relative quantities of the particular polysaccharide groups at different depths of an intervessel PM.

**Figure 4.**
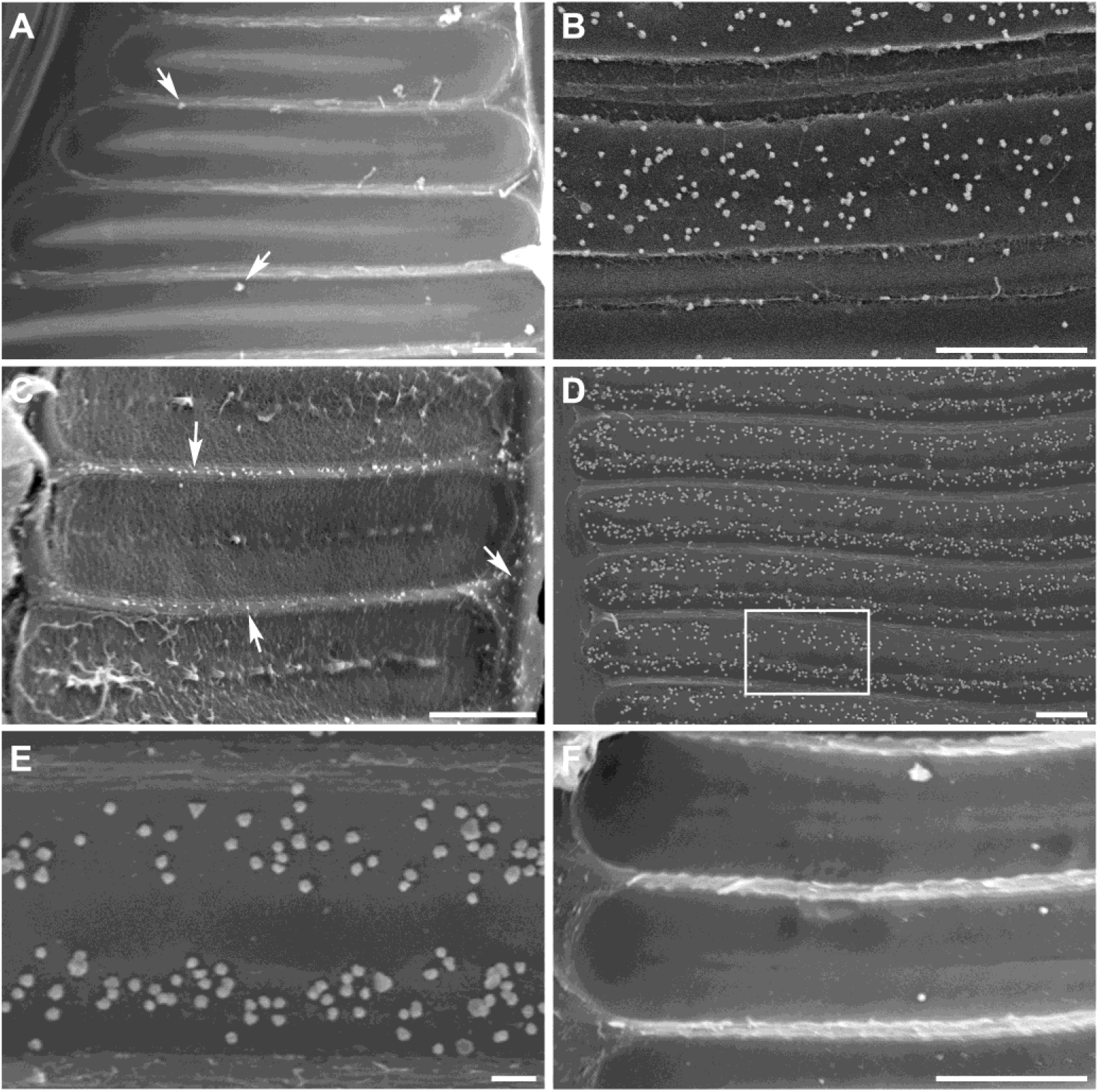
Representative SEM images of the pectic and hemicellulosic polysaccharide detection on Type I intervessel PMs. A, An immunolabeling control (xylem tissue incubated with CCRC-M140 but without its corresponding secondary Ab, and followed by the silver enhancement treatment) showed only several random bright particles (arrows). B-F, Immunogold labeling with CCRC-M1 (B), CCRC-M140 (C), JIM5 (D, E) and JIM7 (F). Bright particles seen on the intervessel PMs are silver-enhanced gold particles, indicating presence of the polysaccharide group recognized by a particular mAb. B, A lot of randomly distributed bright particles (arrows) indicate presence of a large quantity of F-XyGs on the smooth PM surface. C, Absence of xylans on the PM regions revealed by the lack of bright particles despite their rich presence in the secondary wall underlayers and the primary wall strip (arrows) originally concealed under the secondary wall between adjacent PMs. D, Abundant presence of WMe-HGs on intervessel PMs indicated by densely distributed bright particles. E, Enlargement of the framed region in D, indicating that WMe-HGs are not evenly distributed with an abundant, relatively uniform presence in the PM’s peripheral regions and little amount in its narrow central strip region. F, Absence or rare presence of HMe-HGs on the PMs indicated by very few (3) random bright particles. Scale bar equals 2 μm in E and 5 μm in the other panels.

Type I intervessel PMs with the immunogold labelling reveal the four groups of polysaccharides in the surficial layers of an intervessel PM. As shown in Figure 4, the surficial PM layers exhibited some quantitative differences in the four groups of polysaccharides. There were large amounts of F-XyGs present relatively uniformly all over the PM surface (Figure 4, B). Xylans (Figure 4, C) and HMe-HGs (Figure 4, F) were below the detectable level or occurred very little, respectively on the PM surface. WMe-HGs were present in large quantities in the peripheral regions but absent in the narrow central strip region of the PMs (Figure 4, D and E).

Despite some differences in their surface features, intervessel PMs of both Types II and III displayed the same or very similar pattern in the distribution and quantity of each of the four groups of polysaccharides in the underneath layers of an intervessel PM. An abundant amount of F-XyGs occurred over the whole PM region (Figure 5, A) and they appeared to be associated with the PM’s microfibrils (Figure 5, B). This situation remained unchanged no matter whether the PMs contained small pores or not. Compared to F-XyGs, relatively less amounts of xylans (Figure 5, C and D) and HMe-HGs (Figure 5, G and H) were detected in the rough regions of PMs with small pores. The underneath layers of an intervessel PM contained little amounts of WMe-HGs (Figure 5, E and F).

**Figure 5.**
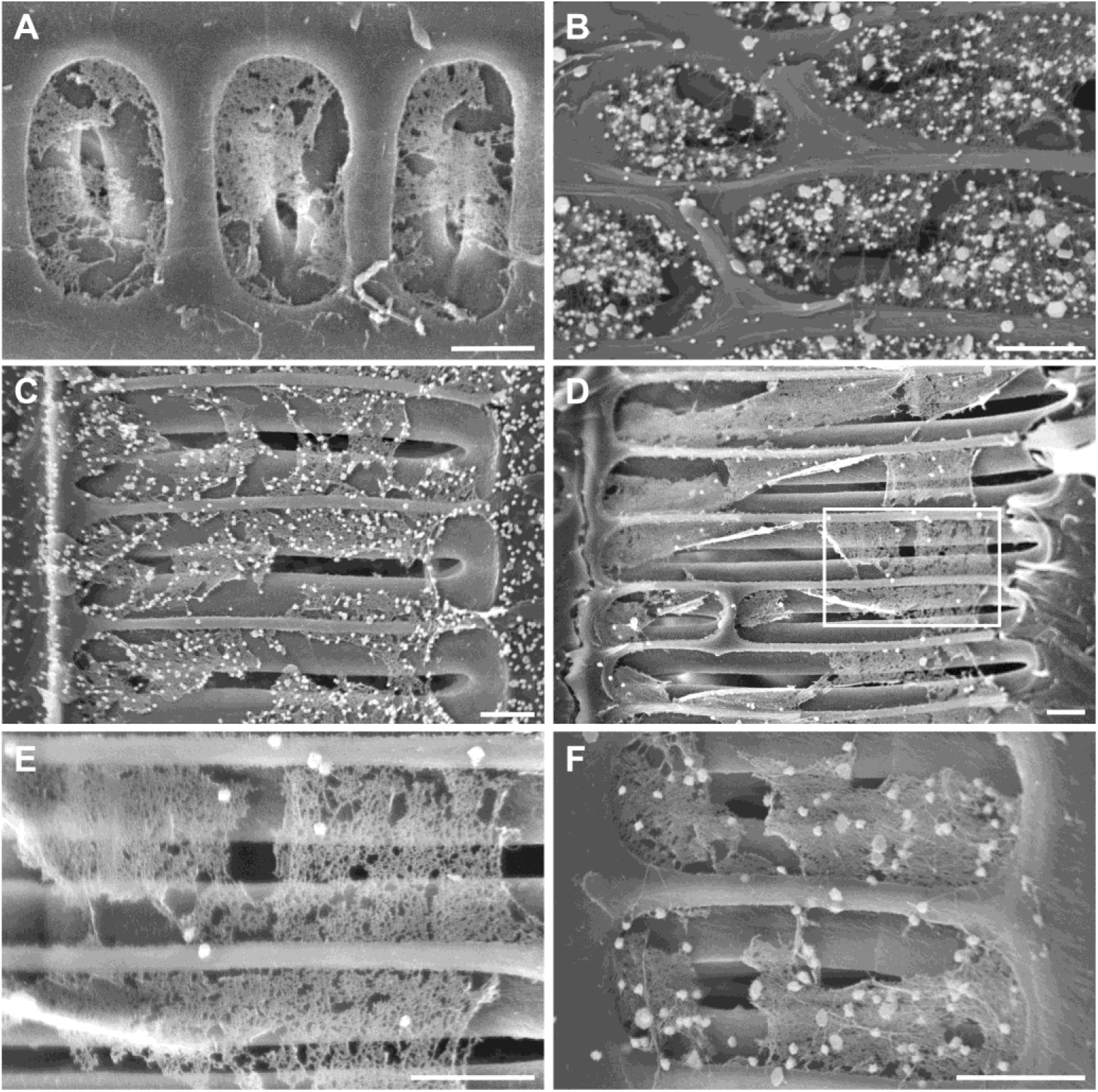
Representative SEM images of the pectic and hemicellulosic polysaccharide detection on intervessel PMs of Types II and III. Intervessel PMs were immunogold-labelled with CCRC-M1 for F-XyGs (A, B), CCRC-M140 for xylans (C, D), JIM5 for WMe-HGs (E, F), and JIM7 for HMe-HGs (G, H). Bright particles indicate the presence of the polysaccharide group recognized by each mAb. A, Dense bright particles indicated an abundant presence of F-XyGs over a whole intervessel PM region. B, Bright particles are associated with microfibrils of the rough PM surface. C, A large quantity of xylans on the PMs revealed by the relative dense distribution of bright particles. D, Bright particles are mostly associated with amorphous material of the PM. E and F, A small quantity of WMe-HGs on the PMs revealed by few randomly distributed bright particles. G and H, A large quantity of HMe-HGs on the PMs indicated by the relatively dense distribution of bright particles. Scale bar equals 5 μm in A, C, E and G, and 1 μm in B, D, F and H.

Type IV intervessel PMs were structurally disintegrated with multiple separate or interconnected highly porous patches of different sizes and shapes, exposing the polysaccharide composition of their deep layers. These highly porous PM regions contained an abundant quantity of F-XyGs (Figure 6, B) and xylans (Figure 6, C), which were uniformly distributed and associated with the PM’s microfibrils. WMe-HGs were present only in a small quantity and randomly occurred to the porous PM networks (Figure 6, D and E). HMe-HGs were present in a large quantity and distributed relatively uniformly over the PM patches (Figure 6, F). Quantitatively, HMe-HGs were still less than F-XyGs and xylans.

**Figure 6.**
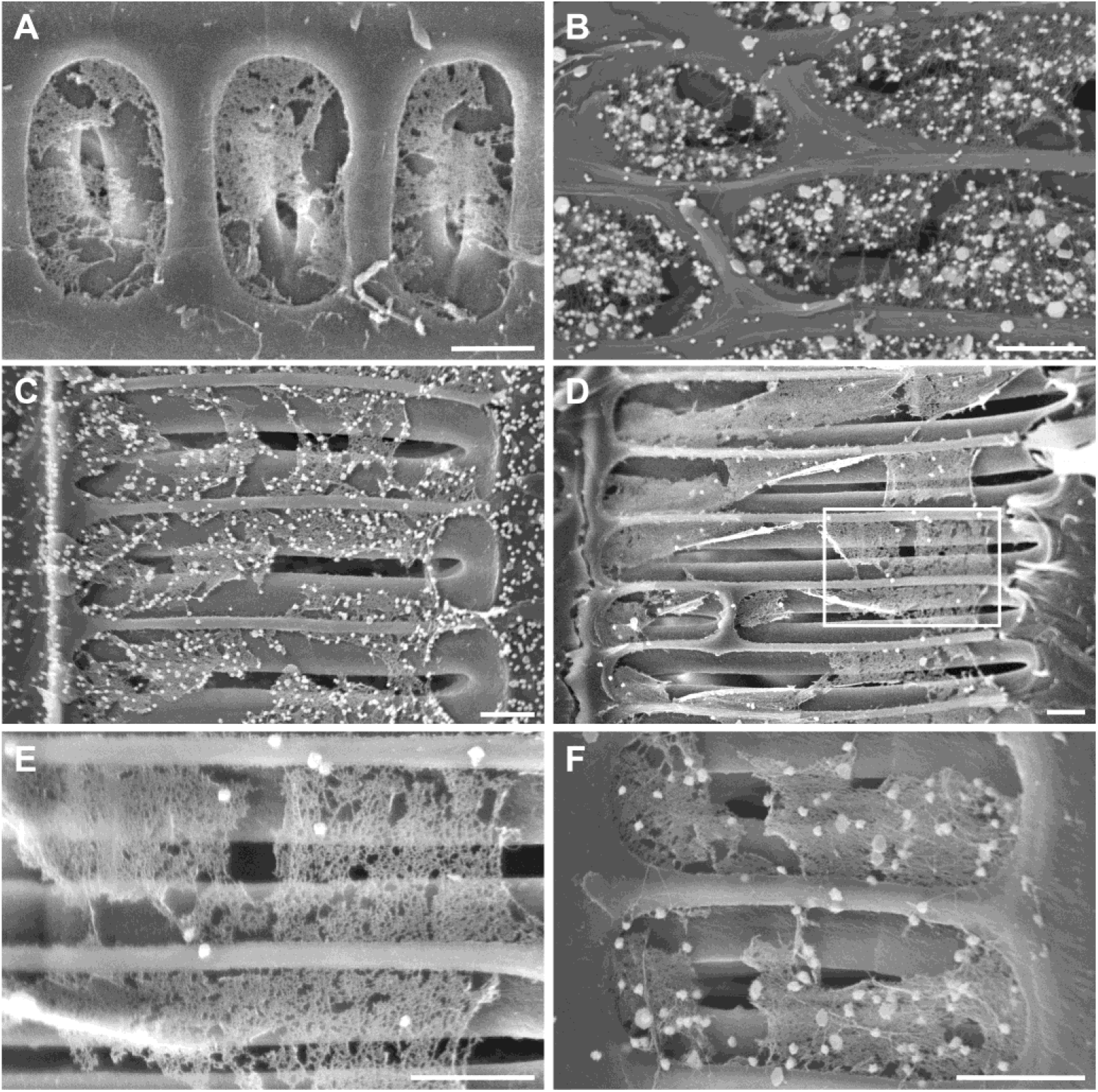
Representative SEM images of the pectic and hemicellulosic polysaccharide detection on Type IV intervessel PMs. A, No bright particles on the highly porous intervessel PMs in an immunolabeling control (xylem tissue was incubated without JIM7 but with its secondary Ab, and then followed by the silver enhancement treatment). Only several bright particles are seen on the highly porous PMs. B-F, Immunogold labeling with CCRC-M1 (B), CCRC-M140 (C), JIM5 (D, E) and JIM7 (F). Different densities of bright particles showed differential quantities of the four polysaccharide groups on the highly porous intervessel PMs. B, The high density of bright particles indicated an abundant presence of F-XyGs on the intervessel PMs. C, An abundant presence of xylans on the PM patches. D, A small amount of WMe-HGs on the PM patches. E, Enlargement of the framed region in D, indicating few randomly distributed bright particles on the highly porous intervessel PM patches. F, A large quantity of HMe-HGs on the two highly porous PMs revealed by relatively dense and uniformly distributed bright particles. Scale bar equals 3 μm in all panels.

The quantities of the four polysaccharide groups described above are summarized in Table 1 for an analysis of their spatial distributions and quantities in an intervessel PM. F-XyGs were in large quantities in the surficial layers of intervessel PM and occurred abundantly at its different depths. Xylans and HMe-HGs were absent or very little in quantity in the surficial layers but had increased quantities towards depths. WMe-HGs were abundant in the surficial layers but present only in a small quantity at different depths.

**Table 1.** Quantitative comparison of four polysaccharide groups at different depths of an intervessel PM

| Depths in Intervessel PM | F-XyGs | Xylans | WMe-HGs | HMe-HGs |
| --- | --- | --- | --- | --- |
| Surficial Layers | ++* | - | ++++ | +/- |
| Underneath Layers | ++++ | ++ | + | ++ |
| Deep Layers | ++++ | ++++ | + | +++ |

\* "+" and "-" indicate presence and absence of a polysaccharide group, respectively. "+/-" indicates presence in little quantity or absence. An increased number of "+" indicates an increased abundance of presence.

## Discussion

### Structural variations of intervessel PMs

Functional roles of intervessel PMs are attributed to their structures and chemical compositions. The current investigation with grapevine as a model plant has indicated that intervessel PMs in a functional secondary xylem displayed significant surface structural variations and were accordingly classified into the four types. An overwhelming majority (~90 %) of intervessel PMs (Types I and II) remained intact with a smooth or relatively smooth surface with certain homogenous features, about 6 % of those (Type III) showed a progressive loss in their surface integrity, and the remaining 4 % of those (Type IV) only with highly porous PM patches in place. Changes in the surface structural features of intervessel PMs from Type I to Type IV have clearly revealed how cell wall material is removed from an intervessel PM during its degradation. This has also indicated that intervessel PMs with diverse structural variations may co-exist in a functional xylem.

These results are different from a long-standing viewpoint on the structure of intervessel PMs, which believes that intervessel PMs should all have a porous structure to facilitate the efficient water transport in xylem. The viewpoint is originated mostly from some early studies on intervessel PMs in primary xylem (e.g., O’Brien and Thimann, 1967; O’Brien, 1970) and secondary xylem (e.g., Schmid and Machado, 1968; Butterfield and Meylan, 1982) of a few monocot and dicot species. According to this viewpoint, porous intervessel PMs should result from the removal of non-cellulosic material from the PMs by hydrolytic enzymes during the vessel element differentiation and may consequently reduce resistance to the across-vessel water flow, facilitating the water transport in xylem.

These early investigations on intervessel PMs mostly utilized LM and TEM. Since intervessel PMs are small and thin-layered structures, LM usually lacks sufficient resolution and sensitivity in revealing structural and chemical compositional details, while TEM technique reveals only the structures on the sectional view of a thin intervessel PM (details on this to be included in the next section). These methodological limitations make it difficult to accurately comprehend intervessel PM structures by LM and/or TEM only. The main technique in the current study involves SEM, which can examine the broad surface of a whole intervessel PM. This technique can not only compare intervessel PMs of different vessels in a large xylem specimen, but also reveal structural details of individual intervessel PMs with resolution comparable to that of TEM (to be further discussed in the next section). The current finding of the diverse surface structural features of intervessel PMs is attributed to the methodological suitability of SEM on this particular structure.

On the other hand, it is believed that intervessel PMs in functional xylem should retain their integrity to maintain the safety of water transport in the xylem (Tyree and Sperry, 1989; Tyree and Zimmermann, 2002; Stroock et al., 2014). Porous intervessel PMs may allow air bubbles originally in very few vessels to spread through the vessel network, making the xylem vulnerable to cavitation that may fail the vessel network’s water transport function (Sperry and Tyree, 1988; Rockwell et al, 2014). Structures of intervessel PMs were also addressed by using explants from a variety of tree species for air seeding experiments, sometimes, along with SEM analysis. Some of these analyses were focused on interspecific differences on intervessel PMs in relation to their differential vulnerabilities to cavitation. It has been found that intervessel PMs may vary in their pore size and thickness among species (Choat et al., 2003; Sano, 2005; Jansen et al., 2007) and thicker PMs often displayed a smoother surface (Jansen et al., 2009). However, the coexistence of multiple types of intervessel PMs with different porosity/structural integrity, thickness and surface features in a same secondary xylem tissue has not been found in record. In addition, the structural variations in the four intervessel PM types reported in this study occur to a much greater extent even than those of intervessel PMs reported among species. Out of the four types, Type IV intervessel PMs, despite the presence in small quantities, displayed the minimal in the structural integrity (Figure 2, G and F; Figure 6). It seems that they well fit the “rare leakier” intervessel PMs category hypothesized in some species more vulnerable to cavitation (Christman et al., 2009; Christman et al., 2012). Therefore, the revelation of the various intervessel PM forms in a same xylem tissue adds a piece of important information that should be carefully considered when analyzing intervessel PM’s impacts on the safety and efficiency of water transport in the xylem.

One issue with SEM in analyzing delicate structures is the possibility of structural artifacts. Any step of sample processing and specimen examination under SEM (i.e., sample trimming, dehydration, sputter coating, and specimen-electron beam interactions) could potentially cause structural artifacts (Carlquist, 1988; Jansen et al., 2008; Plavcová et al., 2011). We saw some structural artifacts with intervessel PMs in some of our early experiments. Intervessel PMs with artificial structures show features very different from what has been observed in the specimens of the current study. With the improved protocol and skills for the sample processing and SEM operating conditions, we have significantly minimized structural artifacts. Also, specimens for our work were processed consistently with the same protocol and many of their specimens were even processed together throughout the whole procedure. The different types of intervessel PMs reported in this study were found in same xylem specimens and/or in vessels close to one another in many cases. This excludes the possibility of the four types of intervessel PMs from being derived from artifacts in the specimen preparation process. As indicated in the Results, the structures of even highly porous PM patches (i.e., Type IV intervessel PMs) can be still well maintained for SEM analysis at high resolution. Therefore, the structural variations of intervessel PMs in this study reflect the actual status of intervessel PMs in the secondary xylem tissues investigated.

It is unclear about what is behind these structural variations of intervessel PMs. Are intervessel PMs with the intact/smooth surface protected from the enzymatic attack during the vessel element differentiation? Lignified primary cell wall and highly methyl-esterified pectins were found to be more resistant to enzymatic degradation (Meylan and Butterfield, 1982; Fromm et al., 2003; Boyce et al., 2004; Kim and Daniel, 2014). The current study and some of our earlier work (Sun et al., 2011) also found the presence of some pectic polysaccharides with different degrees of methyl esterification in intervessel PMs. Another less likely possibility is that intervessel PMs with the smooth surface be derived from sequent modifications on the more porous intervessel PMs resulted from the enzymatic degradation. In some species, parenchyma cells adjacent to a vessel, and developing tyloses in a vessel were found to secrete pectic polysaccharides-rich gels to the vessels (Rioux et al., 1995; Rioux et al., 1998; Soukup and Vitrubová, 2005; Sun et al., 2008). Part of the gels secreted may deposit onto intervessel PMs, displaying a smooth or less porous surface of the PMs. These are all needed to be clarified and will be important to better understand how intervessel PMs can maintain a balance in the efficiency and safety of water transport in a functional xylem.

### Profiling of spatial polysaccharide distribution in intervessel PMs and its importance in analyzing functions of intervessel PMs as well as primary cell wall in general

Some functions of intervessel PMs are affected by the PM’s polysaccharide composition, as suggested by the two major hypotheses about the possible roles of intervessel PM polysaccharides in regulating the PM’s resistance to the water flow in a vessel system (Zwieniecki et al., 2001; Doorn et al., 2011). There have been a lot of investigations on the non-cellulosic polysaccharide compositions of intervessel PMs. An overwhelming majority of them focused on pectic polysaccharides and reported their absence in intervessel PMs (O’Brien, 1970; Butterfield and Meylan, 1982; Catesson, 1983; Wisniewski and Davis, 1995; Jansen et al., 2004; Arend et al., 2008; Choat et al., 2008; Nardini et al., 2011). Some investigations intended to detect hemicellulosic polysaccharides in intervessel PMs similarly reported their absence (O’Brien, 1970; Butterfield and Meylan, 1982; Alves et al., 2009). However, contrary to those reports, the current research has revealed presence of the four major groups of pectic and hemicellulosic polysaccharides (WMe-HGs, HMe-HGs, F-XyGs and xylans) in intervessel PMs.

The different conclusions on polysaccharide composition of intervessel PMs are likely due to the differences in research methods. Detection of polysaccharides in an overwhelming majority of these investigations has been mostly based on some dye staining techniques with LM and/or TEM. They focused on the detection of polysaccharides on a section of intervessel PMs. The effectiveness of a dye staining technique may be determined by multiple factors, including the staining ability of a dye, the dimension/size of a wall structure, and the abundance of target polysaccharides in a structure. Intervessel PMs are very small, thin-layered structures. If the quantity of target polysaccharides is below a certain level, it might not be sensitive enough for the dye to reveal the presence of target polysaccharides. The intrinsic low resolution of LM can further complicate the detection and makes it impossible to reveal any spatial and quantitative differences of target polysaccharides throughout the thickness of an intervessel PM.

In recent years, immunogold-TEM were developed to detect polysaccharides of intervessel PMs in some species (Plavcová and Hacke, 2011; Kim and Daniel, 2013). In this technique, the transverse section of thin intervessel PMs is first treated with a cell wall mAb (primary Ab) that recognizes and binds to a specific polysaccharide/polysaccharide group on the PM. A following treatment with a corresponding nanogold particle-tagged secondary Ab attaches the secondary Ab to the primary Ab. Then, examining nanogold particles under TEM helps localize the target polysaccharides in the transverse section of the PMs. Due to the high sensitivity and specificity of the mAbs, the immunogold-TEM has revealed presence/absence of some pectic and hemicellulosic polysaccharides (Plavcová et al., 2011; Kim and Daniel, 2014, 2016; Herbette et al., 2015). However, just like LM, immunogold-TEM is suitable for detecting polysaccharides mostly in a restricted area on a transverse section of intervessel PM (as shown in Figure 1, C), and is not possible to reveal polysaccharide distribution over a large/the whole surface area of an intervessel PM. These limitations can hinder a comprehensive analysis of polysaccharide components and their spatial distributions and may therefore affect accuracy of the information gained.

The current study used a different technique, immunogold-SEM technique (Sun et al., 2017), to analyze polysaccharide composition of intervessel PMs. This technique also takes advantage of the high sensitivity and specificity of cell wall mAbs in detecting pectic and hemicellulosic polysaccharides but uses SEM to localize target polysaccharides over a large or the whole surface area of cell wall structures. This immunogold-SEM technique has been proved powerful in analyzing polysaccharide compositions of both primary and secondary cell wall (Sun et al., 2017; Zheng et al., 2018). In terms of revealing polysaccharides of intervessel PMs, our immunogold-SEM protocol has three major demonstrated advantages over LM and/or TEM: 1) much higher sensitivity, 2) ability of elucidating polysaccharide composition over a large wall surface area and 3) simultaneous analysis on cell wall structure and its polysaccharide composition up to high resolution. These advantages contribute to the accurate information of the structures and spatial polysaccharide profiling of intervessel PMs.

With the immunogold-SEM, the current study revealed not only the presence of WMe-HGs, HMe-HGs, F-XyGs and xylans but also their differential spatial distribution in an intervessel PM, with abundant WMe-HGs only in the surficial layers, HMe-HGs and xylans more abundant toward depths, and F-XyGs abundantly present throughout the thickness of an intervessel PM. An intervessel PM includes primary cell wall in structure. The differential spatial distributions of these polysaccharides may reflect how these polysaccharides are secreted to a primary cell wall at different stages of the wall deposition. Thus, pectic and hemicellulosic polysaccharides should not be considered to have a homogeneous distribution across the thickness of a cell wall.

On the other hand, multiple structural models of primary cell wall have been established mostly based on analyses of polysaccharide components, particularly hemicellulosic and pectic polysaccharides in the cell wall (Carpita and Gibeaut, 1993; O’Neill and William, 2003; Albersheim et al., 2010; Cosgrove, 2014). It is clear that the polysaccharide composition of a primary wall has a large impact on the wall’s functions. Therefore, accuracy information of polysaccharide components and their spatial distributions becomes important to reveal more details about/better understand cell wall structure and function. The four representative groups of polysaccharides were analyzed in this study; however, other polysaccharides may also be clarified with their specific mAbs by using this model system. Therefore, a further analysis with this system should provide accurate information of the quantity and spatial distribution of different polysaccharides not only in intervessel PMs but also in a primary cell wall. We believe that information gained will be essential for analyzing cell wall formation mechanism, interactions of its polysaccharides, and functional modifications derived from the changes in polysaccharide composition and wall thickness.

### Spatial polysaccharide profiling of intervessel PMs and its importance in analyzing host plant’s disease susceptibility mechanism

In most plant vascular diseases caused by xylem-limited pathogens, disease symptom development largely depends upon pathogens’ spread through the vessel system of host plant (Beckman, 1987; Purcell and Hopkins, 1996; Bove and Garnier, 2002; Newman et al., 2003; Sun et al., 2013). Intervessel PMs are barriers that pathogen must break to achieve its systemic spread in a host plant. It is believed that pathogen secretes CWDEs to attack polysaccharide targets of intervessel PMs and by decomposing the polysaccharides, it may break the barriers for its spread (Roper et al., 2007; Ellis et al., 2010; Sun et al., 2011). Revelation of polysaccharide compositions and their spatial distributions in intervessel PMs can therefore provide key information for an accurate analysis of potential interactions between pathogen’s CWDEs and intervessel PM, which is essential to understand host plants’ disease susceptibility mechanism (Roper et al., 2007; Pérez-Donoso et al., 2010; Sun et al., 2011).

Grapevine Pierce’s disease (PD) is such a representative vascular disease with the causal xylem-limited bacterial pathogen,*Xylella fastidiosa*, spreading throughout a host vine’s vessel system for PD symptom development (Hopkins, 1989; Fry and Milholland, 1990a,b; Krivanek and Walker, 2005; Sun et al., 2007, 2008). The pathogen’s genome has indicated its ability of encoding two groups of CWDEs (i.e., polygalacturonase-PG and endoglucanase-EGase, Simpson et al., 2000), which may degrade hemicellulosic and pectic polysaccharides of intervessel PMs. Porous intervessel PMs were observed in PD-susceptible grapevine genotypes and some vessels with porous intervessel PMs were also associated with *X. fastidiosa* (Sun et al., 2011; Ingel et al., 2019). Since HGs and XyGs are polysaccharides common in the cell walls of dicots (Carpita and Gibeaut, 1993; Freshour et al., 2003; Cosgrove, 2005, 2014), their possible interactions with the pathogen’s CWDEs might be related to the disease susceptibility of a host plant. Our previous study with immunofluorescence LM implied that F-XyGs and WMe-HGs were among potential targets of the *Xylella’s* CWDEs and that the quantity and distribution of these polysaccharides should be related to grapevine susceptibility to PD (Sun et al., 2011).

Recently, polysaccharide analysis on two grapevine genotypes (one is Chardonnay that is also used in the current study) with similar PD susceptibility but differential symptom development progression indicated that the genotype with less amounts of F-XyGs in the surficial intervessel PM layer displayed less porous intervessel PMs and slower symptom development (Ingel et al., 2019). This has further demonstrated the possible impacts of the quantity and spatial distribution of intervessel PM polysaccharides on PD symptom progression.

Data from the current study provide important information to understand potential interactions between *Xylella* and intervessel PMs. It has clearly indicated the heterogeneous spatial distribution of the four groups of polysaccharides in the intervessel PMs of Chardonnay vines (a PD-susceptible grapevine genotype) with abundant WMe-HGs and a large amount of F-XyGs in the surficial layers and other components more abundant toward depths. In a previous investigation on this genotype with PD symptoms (Sun et al., 2011), the initial intervessel PM degradation was found to occur on the PM surface, turning a smooth PM surface to a rough one. This was followed by the exposure of microfibrils of increased amounts and the visibility of pores of increased sizes, eventually resulting in highly porous intervessel PMs or even total disappearance of some PMs. Since this process is correlated with the *X. fastidiosa’*s systemic spread and vessels with porous intervessel PMs are often associated with pathogen cells, this, on one hand, demonstrates the roles of the pathogen’s CWDEs in breaching the integrity of intervessel PMs. On the other hand, it also indicates the breaching of an intervessel PM is due to the sequent removal of wall materials from the surface toward depth of an intervessel PM.

F-XyGs and WMe-HGs are present in large quantities in the surficial layer of intervessel PM. EGase and PG, generated by transgenic *Escherichia coli* that received EGase gene from *X. fastidiosa* and PG gene from *Aspergillus niger,* respectively, were infused to stem explants of the same grapevine genotype to test whether the enzymes may have an impact on its intervessel PMs. It was found that either enzyme alone did not increase the PM porosity, but a mixture of both did (Roper et al., 2007; Pérez-Donoso et al., 2010). Since the two enzymes are known to attack hemicellulosic and pectic polysaccharides. Therefore, based on the data from the current study, it seems plausible to believe that both WMe-HGs and F-XyGs should be at least among the first target polysaccharides of the *X. fastidiosa* CWDEs during the intervessel PM degradation process. This initial degradation may lead to increased porosity in the PM’s surficial layer, making pectic and hemicellulosic polysaccharides in its underlayers more accessible for sequent degradation by the bacterial CWDEs. This may be how xylans and HMe-HGs at depths and other possible polysaccharides present throughout the thickness of an intervessel PM are removed during the PM degradation. Of course, more detailed analyses on this degradation process are still needed to reveal these CWDE-intervessel PM interactions. Analyses from these aspects are essential to clarify the mechanism of pathogen spread in a host vine, consequently helping reveal vascular disease susceptibility mechanism of host plants.

## Materials and Methods

### Plant materials

*Vitis vinifera* var. Chardonnay was used in this study. Experimental vines were grown from commercial rootings in 7.6 L pots with a 16 h light and 8 h dark daily cycle in the Department of Biology Greenhouse at the University of Wisconsin-Stevens Point. Each vine was first trained to retain only two robust buds close to its shoot base, from which two vertical shoots were derived. Each shoot was maintained to have 20 – 25 internodes in height by pruning off its top and some of its lateral branches regularly. When each shoot was 16 – 18 weeks old, a 3 – 4 cm stem length was cut off from the 6^th^ internode (5^th^ if the 6^th^ was surfaced damaged or too short) counting from the shoot base and fixed in 4 % paraformaldehyde in PEM buffer (50 mM PIPES, 5 mM EGTA, 5 mM MgSO4, pH 6.9; Willats et al., 2002) for at least 24 h. A total of 16 internodes from 10 vines were used in this investigation.

### LM, conventional SEM, and TEM for structural analyses of intervessel PPs and PMs

LM, conventional SEM and TEM were used to study structural features of intervessel PPs and PMs. Details of LM were described in Sun et al. (2017). Briefly, multiple 1mm thick xylem tissue block specimens with a surface dimension of 3 mm x 3 mm were cut out from each prefixed sample, exposing a transverse, tangential or radial surface. After washing in 50 mM PIPES, xylem specimens were dehydrated via an ethanol series until 95 % ethanol. Following this was infiltration and embedding of xylem specimens with L. R. White resin (hard grade). Resin blocks were then cured under UV light at – 20°C in a cryochamber (Pelco UVC3, Ted Pella, Inc., USA). 1 μm thick sections were cut with a glass knife from each resin-embedded tissue block on an ultramicrotome (Leica EM UC6, Leica Microsystems GmbH, Austria) and stained with 0.5 % toluidine blue in 0.5 % sodium borate. Stained sections were examined and photographed under a compound light microscope (Nikon Eclipse 50i, Nikon Corp., Japan) equipped with a digital camera (Nikon Digital Sight-5Mc, Nikon Corp., Japan).

TEM was used to reveal structural details of intervessel PP and PMs. Sample fixation and specimen processing were the same as described above for the LM except that specimens were trimmed into cubes of 1 mm side. Each resin embedded specimen was cut into 80 nm-thick sections with a diamond knife (Diatome Ultra 35°, Diatome Ltd., Switzerland) with the same ultramicrotome. Sections were picked up with nickel grid and stained with 2.5 % aqueous uranyl acetate for 15 min. After that, sections were observed and photographed at 75 kV with a TEM (Hitachi 600AB, Hitachi Science Systems, Ltd., Japan).

Samples for conventional SEM were processed following the protocols described in Sun et al. (2006). In brief, secondary xylem specimens from each internode were trimmed and processed for the first few steps in the same way as for the LM. After dehydration with 95 % ethanol, specimens were further dehydrated with 100 % ethanol twice and then critical-point-dried (DCP-1, Denton Vacuum, Inc., USA). Dry specimens were coated with gold-palladium (Desk II, Denton Vacuum, Inc., USA), followed by examination with secondary electron (SE) detector under an SEM (Hitachi S3400N, Hitachi Science Systems, Ltd., Japan) at an accelerating voltage of 8 kV.

Quantitative analyses on the dimension (width and height) of intervessel PMs were based on 246 intervessel PMs on nine samples from six vines. Completely exposed intervessel PMs with the secondary wall borders removed were photographed with SEM and measured for their height and width at the highest height and widest width, respectively. The thickness of intervessel PM was measured on TEM images of 20 intervessel PMs from six samples from four vines. Each intervessel PM appeared to have relatively uniform thickness throughout its width in the transverse section. Frequencies of different intervessel PM types in xylem tissue were based on 54 specimens from five vines (8 – 13 specimens per vine). Three to eight vessels with completely exposed intervessel PMs from each specimen were used in calculating the frequency of each intervessel PM type. Intervessel PMs with features intermediate between two types but closer to one type were counted towards the type for its frequency calculation.

### Immunogold SEM for spatial distribution analysis of hemicellulosic and pectic polysaccharides in intervessel PMs

The current study applied the immunogold-SEM protocol established in a previous work (Sun et al., 2017) to detect multiple hemicellulosic and pectic polysaccharides and their spatial distributions and quantities in intervessel PMs of stem secondary xylem. Four kinds of cell wall mAbs (CCRC-M1, CCRC-M140, JIM5 and JIM7) were used to detect F-XyGs, xylans, WMe-HGs) and HMe-HGs, respectively, in intervessel PMs. All the mAbs were obtained from the CarboSource Services at the University of Georgia (USA). Two kinds of gold-conjugated secondary Abs, goat anti-mouse IgG (whole molecule, batch no. 089K0961) and goat anti-rat IgG (whole molecule, batch no. 129K2049), were used to localize the mAbs of CCRC M and JIM series on intervessel PMs, respectively.

Details of the immunogold-SEM procedure were described in Sun et al., 2017. In brief, for immunolabeling with each mAb, over twenty 1 mm-thick secondary xylem segments were trimmed out of each pre-fixed stem length, exposing their tangential or radial longitudinal surfaces. After wash with 50 mM PIPES and blocking treatment with 3% non-fat milk powder in 0.2 mM phosphate-buffered saline (MP/PBS, pH 7.4) at room temperature (RT), xylem segments were divided into four sets: one for immunolabeling treatment and the others for three types of negative controls. Xylem segments for immunolabeling treatment were incubated in a mAb at 4°C overnight, followed by its corresponding secondary Ab at RT for 1 h. Concentrations of mAbs and secondary Abs were as follows: a 3-fold dilution of the original mAb mouse hybridoma supernatant for each of the two CCRC M Abs, a 10-fold dilution of the original rat hybridoma supernatant of either JIM 5 or JIM 7, a 50-fold dilution of the original secondary Ab for either secondary Ab. Following the Ab incubations were specimen washes first with PBS and then with distilled deionized H2O (DD H2O). Xylem segments were then silver-enhanced at RT for 15 min with a silver enhancement kit (BBI Solutions, UK, Batch No. 12348). After wash with DD H2O and dehydration via an ethanol series, specimens were critical-point-dried and coated with gold-palladium as described above for the conventional SEM. Coated specimens were examined at an accelerating voltage of 8 kV under the same SEM with either a SE detector or a backscattered electron detector.

Three types of negative controls were conducted along with the immunolabelling treatment of each mAb, following the same protocol as its immunolabelling treatment except the Ab incubation steps. The controls included: specimens incubated without both mAb and secondary Ab (specimens were incubated in 3 % MP/PBS at the two steps, the same below), incubated without mAb but with secondary Ab, incubated with mAb but without secondary Ab. Specimens of the controls were processed in the same way for SEM examination.

## Acknowledgements

This work was funded by the US Department of Agriculture-University of California Pierce’s Disease Research Grants Program (Award Number: 2010-266) and the University of Wisconsin-Stevens Point (UWSP) Professional Development Grants. I am grateful to Mimo He, Kevin Thompson and Yuliang Sun for their help in some specimen preparations for the SEM work and to Megan Kitzrow, Ashley Thiel, Kai Chang and John Hardy for their help in taking care of our experimental vines in the UWSP Department of Biology greenhouse.

## Author Contributions

Qiang Sun designed the research, performed research, analyzed data and wrote the manuscript

